# LAVA: a streamlined visualization tool for longitudinal analysis of viral alleles

**DOI:** 10.1101/2019.12.17.879320

**Authors:** Michelle J. Lin, Ryan C. Shean, Negar Makhsous, Alexander L. Greninger

## Abstract

With their small genomes, fast evolutionary rates, and clinical significance, viruses have long been fodder for studies of whole genome evolution. One common need in these studies is the analysis of viral evolution over time through longitudinal sampling. However, there exists no simple tool to automate such analyses. We created a simple command-line visualization tool called LAVA (Longitudinal Analysis of Viral Alleles). LAVA allows dynamic and interactive visualization of viral evolution across the genome and over time. Results are easily shared via a single HTML file that also allows interactive analysis based on read depth and allele frequency. LAVA requires minimal input and runs in minutes for most use cases. LAVA is programmed mainly in Python 3 and is compatible with Mac and Linux machines. LAVA is a user-friendly command-line tool for generating, visualizing, and sharing the results of longitudinal viral genome evolution analysis. Instructions for downloading, installing, and using LAVA can be found at https://github.com/michellejlin/lava.

## Introduction

With the rapid and significant advancements in sequencing technologies in recent years, whole-genome sequencing has become more cost-effective, more efficient, and more accurate than ever (1). A common area of bioinformatics research in virology is the comparison of viral evolution in longitudinal samples. At a basic level, viral genome evolution may be examined over routine passage in cell culture (2,3). Drug manufacturers routinely check for the development of resistance mutations in response to *in vitro* antiviral pressure (4,5). Clinical researchers want to know how viruses evolve longitudinally in normal or immunocompromised patients, in response to a drug pressure, or in different areas of the body (3,6,7).

In order to facilitate these routine analyses of viral evolution, we developed a simple command-line tool called Longitudinal Analysis of Viral Alleles (LAVA) for analyzing and visualizing the evolution of minor variants in viral genomes over time. The basic tenor of these analyses involves the calling of a consensus genome for the initial sample and then using that genome as a reference for downstream samples. Viral sequence data is plotted both across the genome to show where mutations cluster and over time to show allele frequency changes. The metadata associated with the experiment may be minimal, consisting simply of sample names and units of time. The units of time are arbitrary and may be minutes, hours, days, months, years or even different categorical experimental conditions. LAVA also generates interactive HTML files for sequence data analysis. The HTML files may be manipulated by users without significant bioinformatic experience according to the nature of their biological question, alleviating a significant conundrum for sequencing and bioinformatics groups as demand for their services continues to increase.

## Methods

LAVA (Longitudinal Analysis of Viral Alleles) can be downloaded at https://github.com/michellejlin/lava. Installation and usage instructions, a folder with example inputs, and the full source code, are also available at this link. The general workflow is shown in Figure 1. A brief explanation is also given here, but a more in depth look into the pipeline is available at the GitHub link, including options and arguments passed to third party tools, the full LAVA source code, as well as an informative readme document.

**Figure 1.**
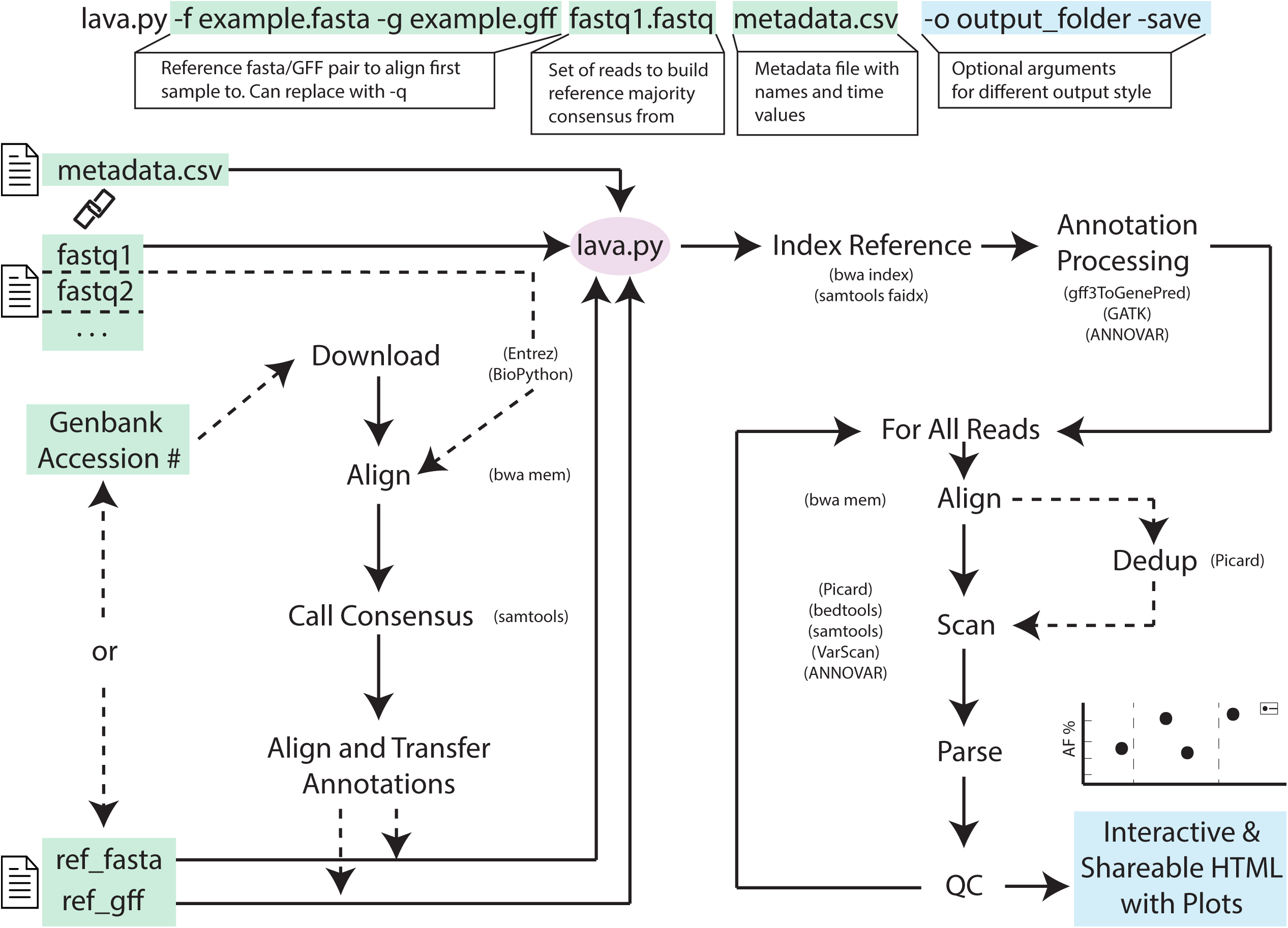
General workflow of the LAVA pipeline is depicted to offer a high-level overview of program execution. Dashed arrows represent optional steps. Input are shown boxed in green, output in blue, and the main lava program is circled in pink. For input, either a GenBank Accession number or a FASTA/GFF pair is required. If a GenBank Accession number is provided, LAVA generates a FASTA/GFF pair following the outlined steps. The linked chain symbol between the metadata.csv input and the FASTQ reads is meant to emphasize that the metadata.csv must contain all the file names that you wish to include in your analysis. General steps are given with tools used during that specific step listed to the side or underneath each step in parentheses. The final output is given as HTML files that contain the interactive plots. For exactly what is passed to each of the other programs and information about parameters and optional arguments (such as mapping parameters), the source code is available on GitHub.

Installation of LAVA and all required dependencies is performed by an install script which is included in the GitHub repository. The install script only requires Python, a Java runtime environment, brew/apt-get for Mac/Linux systems, and an Internet connection. All third party tools except for ANNOVAR (8), which must be manually registered for and downloaded, are also automatically installed. The install script can also be run in ‘check mode’ and the script will check for all required dependencies and print error messages with instructions for how to fix any missing dependencies. The GitHub readme also contains a walkthrough for manually installing all dependencies and LAVA.

Before execution, LAVA requires a reference genome for sequencing read alignment, which can be provided as either an NCBI GenBank Accession number, or a local nucleotide FASTA file along with a GFF file to provide gene and protein annotations (9). Sequencing reads for all samples are input as adapter and quality-trimmed FASTQ files. LAVA currently does not perform adapter or quality trimming so this needs to be done beforehand as required. A two-column CSV metadata file is also required to match sample with its longitudinal temporal information such as passage number or day number. Other available options include removing PCR duplicates for metagenomic sequencing, manually specifying the name of the output folder, analyzing variants by nucleotide changes instead of amino acid changes, saving all intermediate files (which LAVA by default removes), automatically specifying an allele frequency cutoff (which is by default set at 1%), and saving output as PNG files.

There are two different methods of selecting annotations for the reference sequence: automatic and manual. In automatic mode LAVA begins with searching GenBank for a user provided accession number corresponding to the viral species to be analyzed (10). This record is then downloaded both as a nucleotide FASTA file and the complete GenBank record. LAVA aligns the first FASTQ file provided to the downloaded reference (11,12) and calls a majority consensus sequence based off this alignment using samtools (13–15). Then coding sequence annotations are pulled from the GenBank record and transferred to the new majority consensus FASTA using a MAFFT alignment (16). In manual mode the user specifies a reference FASTA and a GFF file containing protein annotations for this reference sequence. LAVA assumes that the FASTA is the majority consensus for the first sample and the GFF is a correct annotation of the reference nucleotide sequence. The result of both the automatic and manual processes is an annotated majority consensus of the first sample.

LAVA then aligns each of the FASTQ files specified in the metadata CSV to this newly generated file, using bwa-mem (17). By default, LAVA does not remove PCR duplicates given the common use of AmpliSeq-like approaches for viral genome sequencing; however, this option can be added in cases where removing PCR duplicates would give a more accurate representation of the data, such as the analysis of metagenomic samples. Variants are called for every position in the genome for every sample using VarScan and saved as standard VCF files (18). These files are removed from the output folder during cleanup to keep disk usage low, but can be saved using the --save option. Variants for all bases are annotated using Annovar and GATK as nonsynonymous, synonymous, complex, stop-gain, or stop-loss, along with the coverage and allele percentage at each base (8,19). The main text file generated by the pipeline is a table called merged.csv containing all the samples, their metadata, and all the amino acid changes. This file, along with reads.csv, the individual .bam files, and the individual .genomecov files, is used for generating the interactive visualization but can also be manually parsed and examined for more in-depth or non-standard analysis. Reads.csv provides read mapping information for each sample, such as total number of reads in sample and percentage of reads mapping to reference. A .bam file is generated for each sample during the alignment process, and these can be viewed for understanding the alignments and how the reads were mapped. Genome coverage for each base in each sample is parsed and extracted into a file with extension .genomecov, so genome-wide depth can be examined and analyzed.

LAVA then visualizes this information with the Bokeh Python module (20–22), allowing for an easily readable and interactive data visualization. The output for this step is an HTML file containing two interactive plots (Fig 2). The first plot depicts allele frequency changes for each variant across the genome for each sample. Tabs at the top of the plot allow easy switching between samples. Sliders to the right of the main plot allow the user to dynamically change the visibility of variants by depth and by coverage. To the right side of the plot, a line graph shows per-base coverage across the whole genome to help inform the user of reasonable coverage thresholds. The second plot shows allele frequency across the samples over time. Given that the samples will be representing different time points (passages, cultures, days past infection, etc.) of a single virus, this plot shows the longitudinal evolution of amino acid changes, separated by protein. Tabs at the top of the plot allow the user to specify which protein they want to examine. Allele frequencies for all changes in the selected protein are plotted over time. Here, variants can also be filtered by depth and coverage. Both plots support zooming and panning and each mutational change has an associated tooltip which can be viewed by hovering with the mouse over the associated data point to display locus-associated metadata. Data can be filtered by type of mutation (synonymous, non-synonymous, stopgains/stoplosses, and complex mutations), as well as if the same mutation occurs across multiple samples. Another available option in the command line is to show nucleotide changes such as transversions and transitions rather than amino acid changes, which may be relevant to cases when examining nucleoside-analog antivirals directed against viral polymerases, base editors such as APOBEC or ADAR proteins, or other aspects of viral epigenetics (23–25).

**Figure 2.**
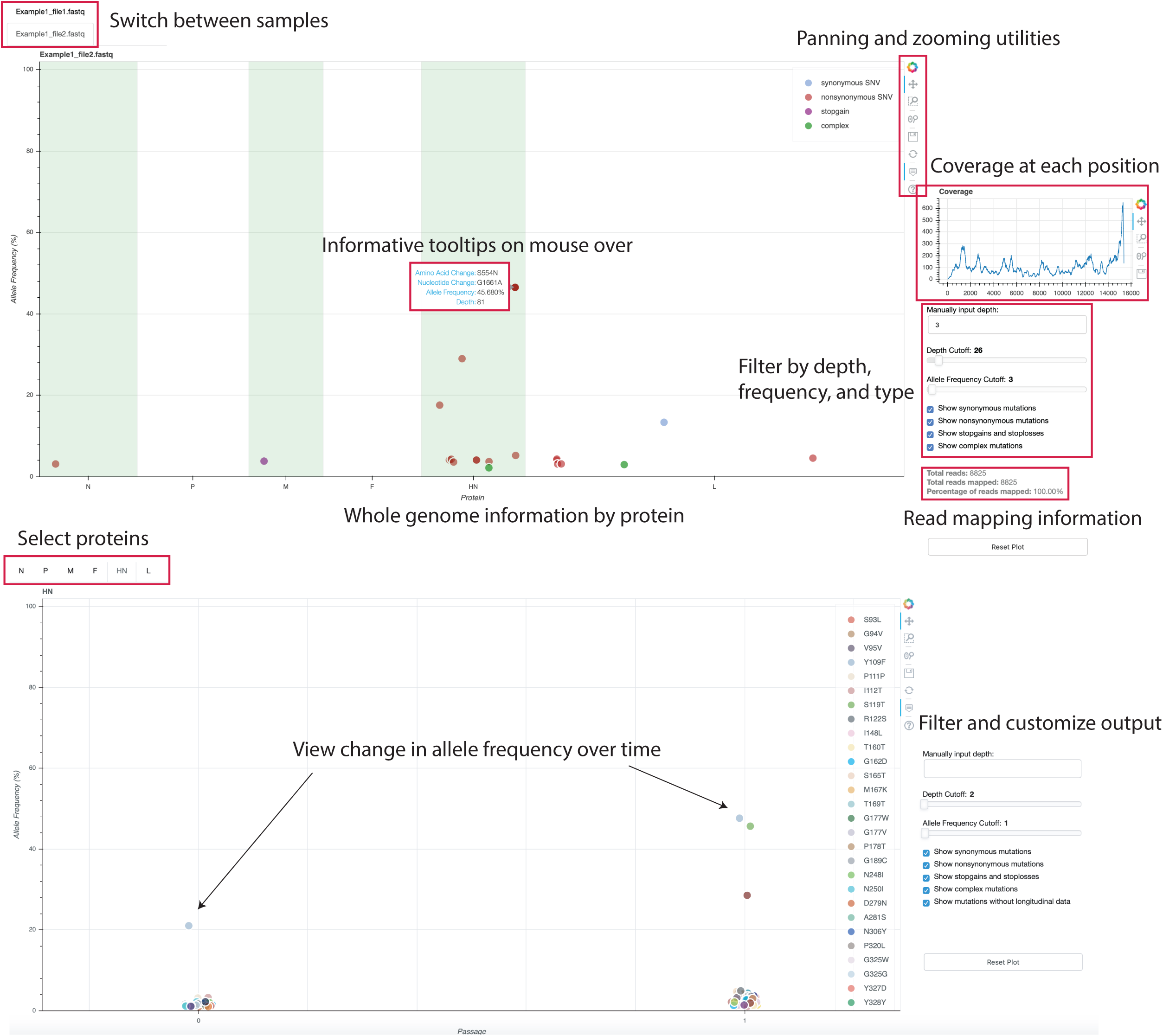
Example LAVA output is shown, this figure shows the results from running Example 1 (All files are available on GitHub and a more in-depth coverage of this data is provided in *Case study 1*.) This example is a screenshot of a Chrome browser displaying the final HTML created by LAVA. The plot on the top of the page shows all amino acid changes across the whole genome for each sample. You can switch between the samples using the tabs highlighted in a red box. The bottom plot shows changes in by-protein allele frequencies over time. You can use tabs once again to switch between proteins. All changes meeting display requirements are plotted over time (or whatever your numerical metadata was). For example, this example shows the hemagglutinin-neuraminidase protein for HPIV3 undergoing changes during the culturing process. All output can be filtered by depth, allele frequency and type of mutation using the sliders boxed in red to the right of each main plot. A small plot is displayed next to the whole genome graph providing a visual representation of the per-base coverage of reads mapping to the consensus.

LAVA outputs all these files in HTML format (Fig 2), which are readily interpretable in any web browser by groups without significant bioinformatics experience. Once generated by LAVA, all these graphs can be sent and shared as standalone files. Additionally, LAVA also has an option to generate static PNG images of the results for situations where interactive visualization is not appropriate such as publications or presentations.

## Results and Discussion

To demonstrate the intended use cases of LAVA and demonstrate why it represents a new and useful tool, we illustrate two real world examples from our own lab.

### Case Study 1 – Evolution of human parainfluenza virus 3 in culture

The provided examples (https://github.com/michellejlin/lava/tree/master/example), which are included with the software, illustrate the automation of a task which the authors first performed manually. For case study 1, these example files are truncated versions of the real data analyzed in Iketani et al. (26), and are named Example 1 in LAVA. Example 1 illustrates how to use LAVA to rapidly perform whole genome analysis on matched samples to understand how a unique selective pressure (i.e. culture exposure) affects viral evolution.

Briefly, paired human parainfluenza virus type 3 (HPIV3) samples were sequenced directly from nasal sampling and after isolation in culture. The study aimed to examine how HPIV3 adapts to brief exposure to culture. Sequencing reads were adapter and quality trimmed using cutadapt, producing the FASTQ files available on the GitHub (27). A simple metadata sheet called Example1_metadata.csv was created containing file names and ‘passage numbers’. In this case, because we only use two samples, we put the first sample (nasal swab SC332) occurring at passage 0 and the second sample (cultured CUL 332) at passage 1. We have also provided a manually generated GFF/FASTA reference pair containing the protein locations for all HPIV3 proteins except C and D (which are created through RNA editing and thus do not automatically translate correctly). To reduce the file size, the example files uploaded to GitHub contain only the first 20,000 original reads that correctly mapped to HPIV3 – full sequencing read files are available from BioProject PRJNA338014. Example 1 shows how rapid adaptations to culture can be discovered using LAVA as two non-synonymous mutations (S554G and P241L) appear in the sample after very brief growth in culture. This example also shows off the utility of the depth and allele frequency sliders which can be used to quickly filter low-level sequencing artifacts and mapping errors out of the data, allowing the user to focus on the most relevant points of data.

### Case Study 2

We have also included data for a case study which fully highlights the longitudinal analysis nature of LAVA. In this study, norovirus samples were recovered from a >250 day infection over 11 time points from a single patient (28). The fundamental question in this analysis is what whole genome changes accrue as norovirus adapts to the immunocompromised host over almost a yearlong period.

Samples were sequenced and reads were adapter and quality trimmed using cutadapt as part of our routine metagenomics analysis pipeline (27). As in Example 1, we selected a few samples: ST107, ST283, and ST709 (all available on BioProject PRJNA338014). Reads were trimmed to reduce file size to upload onto GitHub. A two-column metadata sheet called Example2_metadata.csv was created mapping samples to collection day. The analysis was run with the one-line command “lava.py -q MH260507 ST107.fastq Example2_metadata.csv -o norovirus_output” (MH260507 is the GenBank Accession number for the actual day 0 consensus of these samples). This command showcases the alternative method of generating reference files: using the -q flag to automatically download a GenBank reference and transfer annotations. This example also highlights the utility of the protein plots, which show how the allele frequency of all variants for each protein changes over time. Instead of using passages as in Example 1, these plots demonstrate the evolution over number of days of infection. Using these plots, one can see how the entire norovirus genome accumulates fixed mutational changes over a long-term infection with an increased rate of fixed mutational changes in VP1, the capsid protein and main antigenic determinant of norovirus (29).

### Comparisons

While there are many programs that process and visualize somatic mutations, LAVA is unique in its focus on monitoring minor variant alleles in viruses (30–32). With both its component parts of pipeline and visualizer, LAVA fills an important need in the viral bioinformatics community. The Broad Institute, for example, has several well-documented workflows for both germline and somatic variant discovery: HaplotypeCaller and MuTect2. These tools are excellent for their intended use cases and LAVA uses a workflow inspired by these tools. However, HaplotypeCaller is not well suited for whole genome analysis of viral genomes, as the tool is focused on germline SNPs and does not handle the extreme allelic variance found in viral genomes. MuTect2, the Broad Institute’s somatic SNP and indel caller, performs well for its intended use but does not emit all bases of a genome, which is vital information for viral whole genome analysis. Both of these tools are excellent for their intended purposes but would have to be significantly modified to reproduce the analysis of LAVA.

The Broad Institute’s viral-ngs suite, pipelines designed specifically for the analysis of viral genomics, takes paired-end reads and calls intrahost variants (iSNVs). Taxonomic read identification is also visualized with Krona. For variant calling in viral genomes, viral-ngs is an excellent tool and we recommend using it over LAVA. However, LAVA was created specifically to automatically compare longitudinal data, which is not a built-in feature of viral-ngs. LAVA also has a visualization tool to easily see and compare minor allele variants across the genome and across time. In these use cases, LAVA adds functionality over other bioinformatics programs.

Two other bioinformatics pipelines exist that perform similar tasks as LAVA. SMuPFi is a pipeline that, like LAVA, analyzes NGS data to provide a graphical representation of SNPs and works well for viral analysis (33). However, due to its nature as a tool designed to better understand viral escape mechanisms, SMuPFi operates in the area of co-occurring mutations, and works best with only two co-occurring mutations at the same time due to the complex statistical analysis involved.

Another pipeline that serves to identify variant sites is ViVan (34). ViVan takes similar input as LAVA and has a very easy to use, albeit size limit restricted, web interface. It also detects more sensitive variant alleles than LAVA does—it claims to identify variant alleles with a frequency of >0.1%, with a slightly higher rate of false positives, whereas LAVA by default both filters out any minor allele variants below 1% frequency (though this can be adjusted using the – af argument), and allows dynamic filtration in its visualization to suit the user’s purpose. ViVan searches for variants within each sample individually and currently provides no built-in feature for comparisons between samples.

LAVA combines many of the gold-standard bioinformatics tools into a single pipeline to annotate minor allele variants in viruses and adds a truly unique functionality with its interactive visualization. The plots that LAVA outputs present easily understandable comparisons between longitudinal samples, illustrating complex relationships in a simple format that makes patterns like evolution of minor allele variants across samples, nucleotide change frequency in different proteins, and synonymous vs. nonsynonymous mutations in the genome evident. By allowing dynamic filtering of data by allele frequency and coverage depth, these plots can be adjusted to suit the individual needs of the user.

Additionally, the inherently shareable nature of the HTML plots that LAVA creates as output is an advantage. The small size, ability to be viewed on any web browser, and lack of dependencies allow data to be shared quickly and extensively through email or any other means, especially with collaborators who are not comfortable filtering BAM and VCF files.

### Limitations

LAVA is a powerful tool for analyzing a diverse variety of viral datasets, yet it is not without its limitations. While stopgains and stoplosses are handled correctly and included in the plots, LAVA is currently unable to handle complex mutations, wherein two neighboring nucleotide variants occur within a single codon. Multiple nucleotide changes within the same codon are each treated individually as separate amino acid changes. However, LAVA automatically detects this situation, and both prints a warning to the console and colors points corresponding to complex mutations distinctly. Sequence variations such as copy number changes, recombination, or large deletions and insertions that escape the bwa-mem aligner may also be missed (35). Due to the nature of its visualization, LAVA also does not display overlapping genes properly and instead shows them side-by-side. However, LAVA does print a warning message to the console if overlapping proteins are detected, directing users to the README which contains directions for how to manually prepare a GFF file without overlapping proteins. LAVA also does not correctly analyze proteins with RNA editing or ribosomal slippage. Many of these limitations can be fixed by editing the GFF file accordingly.

Another limitation of LAVA is that web browsers can fail to render the output plots if there are an extremely large number of variants (>5,000). This does not impact the actual analysis, only the visualization, and the merged.csv output file will still contain all relevant data. This could create problems if LAVA was used to analyze bacterial genomes or other extremely large genomes. LAVA will print a warning message if there are greater than 5,000 variants. The nature of the merged.csv output file is such that manual analysis could easily be performed in an environment better suited to visualizing extremely large data sets such as R.

## Conclusions

LAVA allows users to go from sequencing data to dynamically interactive plots illustrating longitudinal changes in their samples. The only required inputs are 1) FASTQ files with sequences for analysis, 2) either a GFF file and reference FASTA or a Genbank accession number, and 3) a simple metadata.csv file containing information about sample name and passage number. LAVA cuts down the time and effort significantly for data analysis of longitudinal samples, and provides an intuitive and interactive visualization that can be easily shared among collaborators.

### Web resources

LAVA can be found at https://github.com/michellejlin/lava and is programmed in Python.

## Acknowledgements

The authors would like to acknowledge the Broad institute for Picard, as well as the developers and maintainers of UCSC Genome Browser for gff3ToGenePred. We would also like to thank the entire open source bioinformatics community for their commitment to producing freely available and useful tools for everyone.

